# *Bordetella* BcrH1 and BcrH2 are specific chaperones for the pore-forming complex

**DOI:** 10.1101/2025.03.27.645768

**Authors:** Toshinobu Ogawa, Yuya Kishino, Hiroko Sato, Akio Abe, Asaomi Kuwae

**Author notes:** T. Ogawa and Y. Kishino contributed equally to this work.

## Abstract

*Bordetella* has a type III secretion system that secretes virulence proteins crucial to the establishment of infection. The genes encoding components of the *Bordetella* type III secretion system are located in the *bsc* region on the chromosome. This region includes the *bcrH1* and *bcrH2* genes, which respectively encode the proteins BcrH1 and BcrH2. In this study, we analyzed the roles of BcrH1 and BcrH2 in *Bordetella* infection. First, we created a BcrH1/BcrH2-double deficient strains, and analyzed the amounts of the type III secreted proteins BopB and BopD, which make a complex that forms pores in the host membrane, in bacterial cells of each protein-deficient strain. The results showed that the BopB and BopD signals were weakened in the whole cell fraction of the BcrH1/BcrH2-double deficient strains, respectively. The hemolytic activity and cell toxicity of each BcrH protein-deficient strain were significantly lower than those of the wild-type strain. When anti-BcrH1 and anti-BcrH2 antibodies were used for the immunoprecipitation assay, the BopB and BopD signals were detected in the precipitated fractions, respectively. These results strongly suggest that BcrH1 and BcrH2 are specific chaperones for maintaining the stability of BopB and BopD, respectively.

## Introduction

*Bordetella pertussis* infects the respiratory tract and causes whooping cough in humans (1). It has been suggested that the type III secretion system plays an important role in the *Bordetella* colonization of the respiratory tract (2). The type III secretion system is conserved in many Gram-negative pathogens, and the pathogenicity of these bacteria is often significantly reduced when the type III secretion system function is lost (3).

The type III secretion system has a needle-like structure that penetrates the inner and outer membranes of the bacteria. These pathogens directly inject a group of proteins called effectors produced within the bacteria into the cytoplasm of mammalian cells. The effectors modulate the signal transduction pathway of the mammalian cells through interactions with host factors to establish bacterial infection (4, 5). In *Bordetella*, an effector called BteA is transferred into mammalian cells via the type III secretion system and induces cell death accompanied by membrane destruction (6, 7).

Through the type III secretion system, the bacteria secrete proteins that form pores in the mammalian cell membrane. The amino acid sequences of these pore-forming factors are homologous among bacteria producing type III secretion systems. In *Shigella*, IpaB and IpaC (8), and in *Yersinia*, YopB and YopD (9), function as pore-forming factors. In *Bordetella*, BopB and BopD have been shown to make a pore-forming complex (10). Pore-forming factors bind to proteins called chaperones to maintain stability within the bacterial cell (11). For example, YopB and YopD maintain stability within the bacterial cell by binding to a protein called SycD (12). A protein called BcrH2 in *Bordetella* bacteria has also been shown to bind to a complex of BopB and BopD, but it has not determined whether BcrH2 contributes to the stability of BopB and BopD within the bacterial cell.

In this study, we attempted to clarify the functions of BcrH1 and BcrH2, and whether these BcrH proteins are associated with the stability of BopB and BopD within the *Bordetella* cytoplasm.

## Results

### The amounts of BopB and BopD are reduced in BcrH1-deficient and BcrH2-deficient strains, respectively

To analyze the function of BcrH1, we first created *B. bronchiseptica* strains lacking BcrH1. We cultured the wild-type, BcrH1-deficient (ΔBcrH1), and BcrH2-deficient (ΔBcrH2) strains, and prepared whole cell fractions and supernatant fractions. We then examined these samples by SDS-PAGE followed by Western blotting. No difference was observed in the BopD signal intensity between the wild-type and ΔBcrH1 strains (Fig. 1), but the BopB signal intensity in the ΔBcrH1 strain was significantly weaker than that of the wild-type strain (Fig. 1B). The BopB signal was detected in the whole cell solution and supernatant fractions of the wild-type and ΔBcrH2 strains with almost the same intensity (Fig. 2), whereas the BopD signal was significantly weaker in the ΔBcrH2 strain compared to the wild-type strain (Fig. 2B). These results suggest that the presence of BcrH1 and BcrH2 is necessary to maintain BopB and BopD, respectively.

**Fig. 1.**
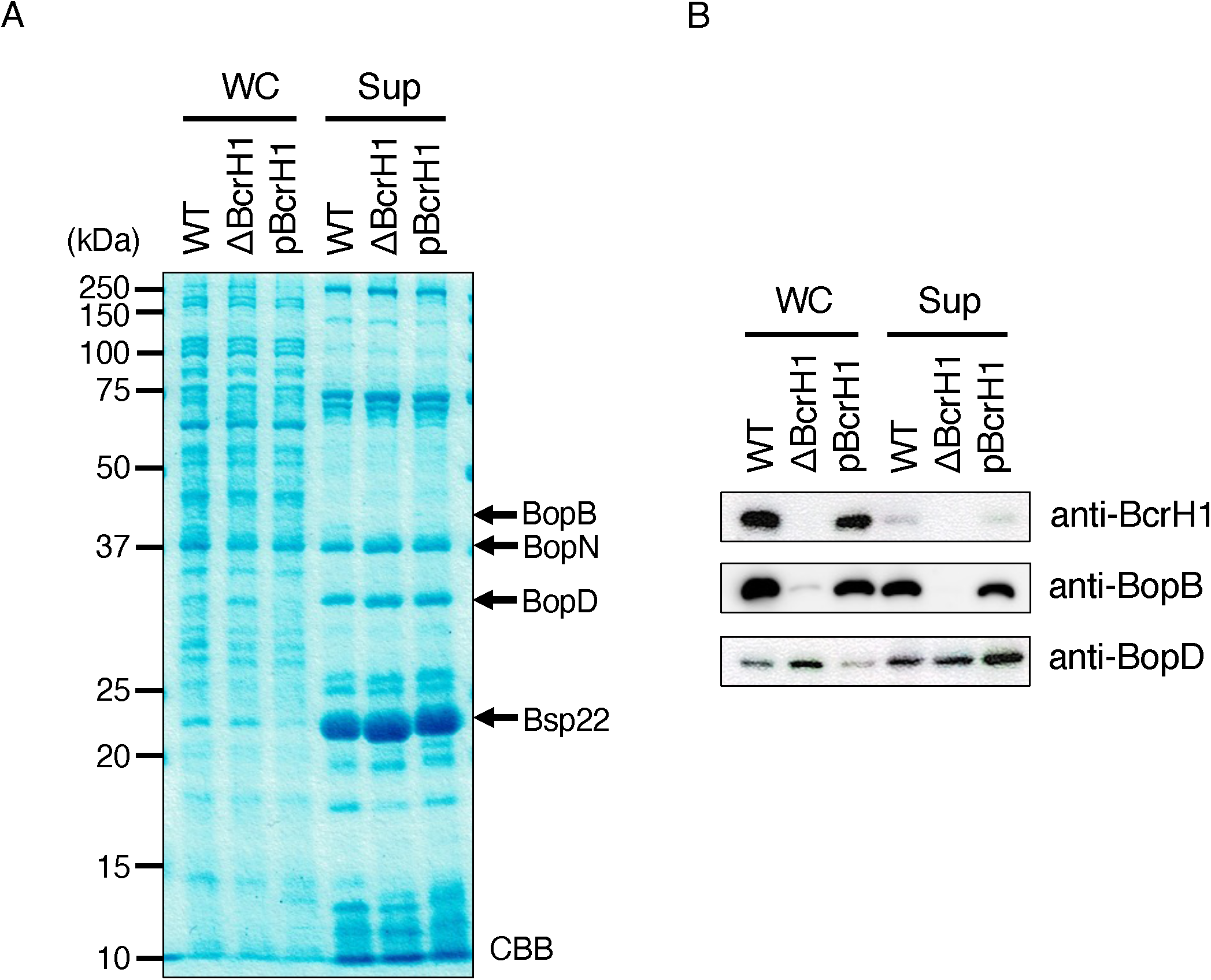
Production and secretion of type III secreted proteins in BcrH1-deficient mutants. The wild-type strain, the BcrH1-deficient strain (ΔBcrH1), and the strain in which pBcrH1 was introduced into ΔBcrH1 (pBcrH1) were cultured in SS medium, and then the whole cell fraction (WC) and the supernatant fraction (Sup) were prepared. (A) The gel shows SDS-PAGE gel stained with CBB after separation of WC and Sup samples by a 12% gel. The numbers indicate the molecular weight markers in kDa. (B) The results of Western blotting using anti-BcrH1, anti-BopB, and anti-BopD antibodies are shown.

**Fig. 2.**
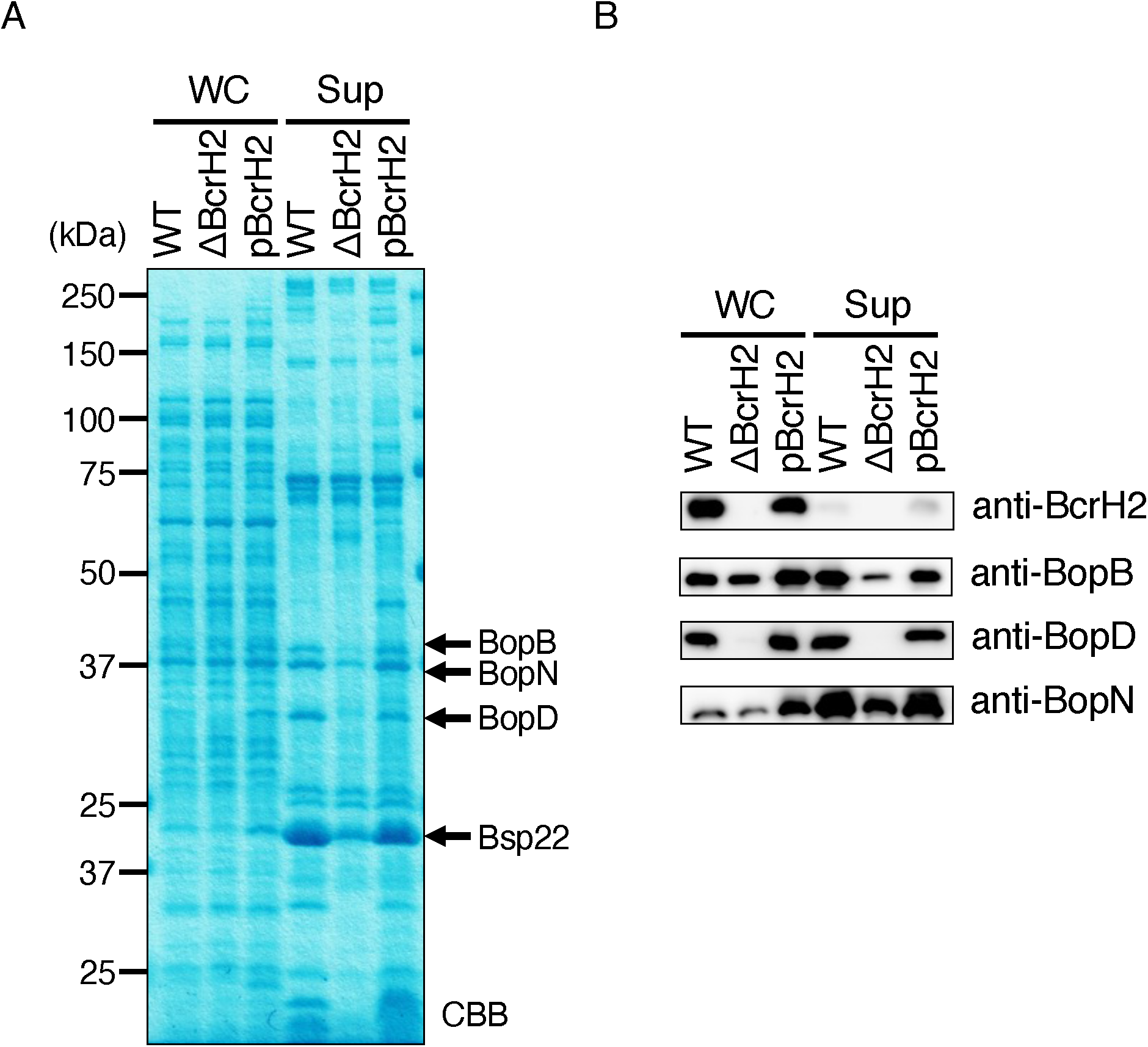
Analysis of the production and secretion of type III secretory proteins in BcrH2-deficient strains. The wild-type strain, the BcrH2-deficient strain (ΔBcrH2), and the strain in which pBcrH2 was introduced into ΔBcrH2 (pBcrH2) were cultured in SS medium, and the whole cell fraction (WC) and the supernatant fraction (Sup) were prepared. (A) A 12% SDS-PAGE gel stained with CBB after separation of the WC and Sup samples. The numbers indicate the molecular weight markers in kDa. (B) The results of Western blotting using anti-BcrH2, anti-BopB, anti-BopD, and anti-BopN antibodies are shown.

### BcrH protein-deficient strains have reduced hemolytic and cytotoxic activities

*Bordetella* destroys erythrocyte membranes in a BopB- and BopD-dependent manner (10, 13). To examine whether the BcrH protein-deficient strains have hemolytic activity, the wild-type strain and the two BcrH protein-deficient strains were cultured and exposed to rabbit erythrocytes. As a result, the hemolytic activities of the BcrH proteins-deficient strains were significantly reduced compared to the wild-type strain (Fig. 3).

**Fig. 3.**
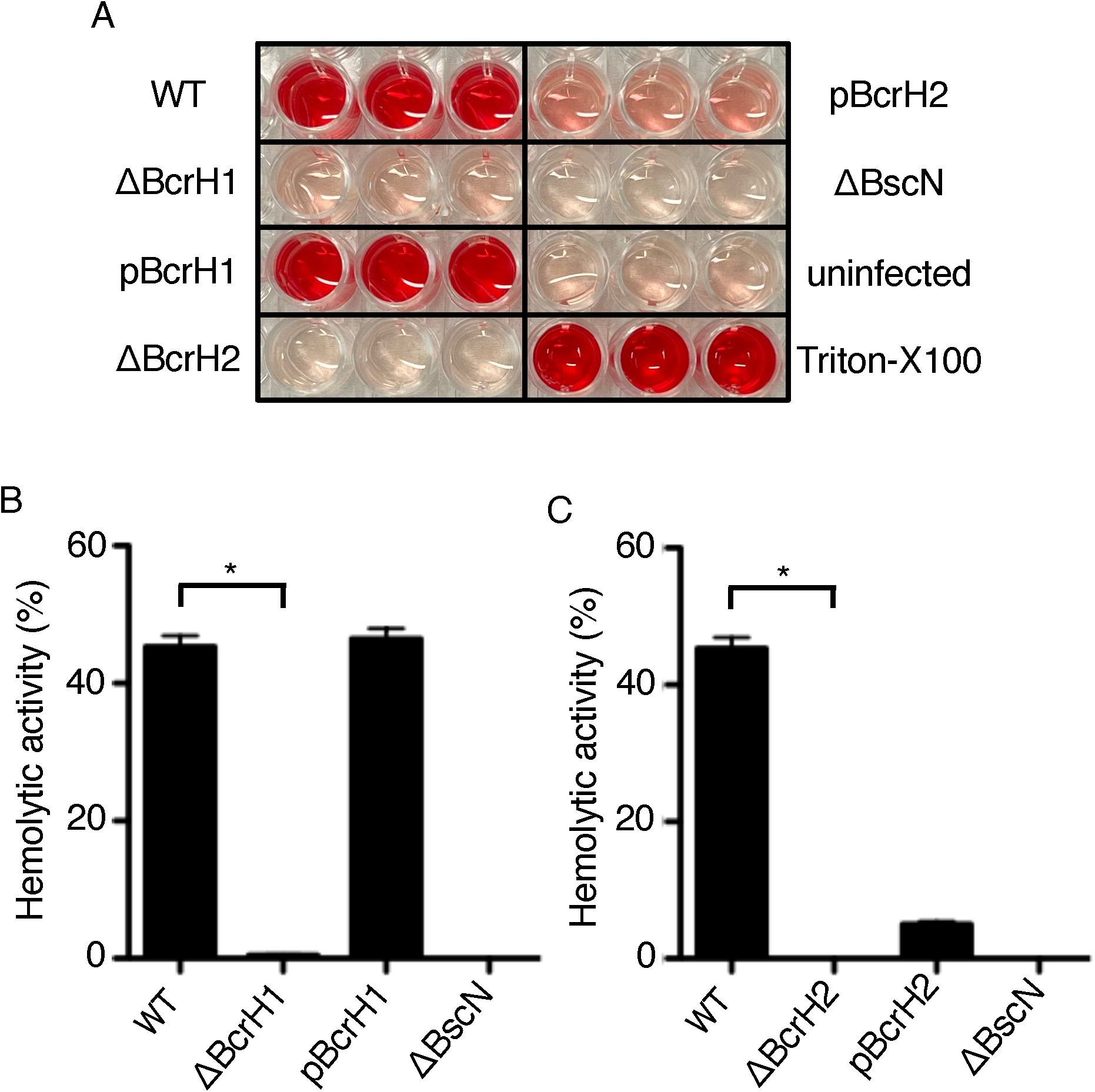
Analysis of the hemolytic activity of BcrH1- and BcrH2-deficient strains. (A) A photograph of a 96-well plate onto which each strain was cultured with rabbit red blood cells for 1 h and the supernatant was transferred after centrifugation. The following *B. bronchiseptica* strains were analyzed: the wild-type strain (WT); the BcrH1-deficient strain (ΔBcrH1); the strain in which pBcrH1 was introduced into ΔBcrH1 (pBcrH1); the BcrH2-deficient strain (ΔBcrH2); the strain in which pBcrH2 was introduced into ΔBcrH2 (pBcrH2); and the type III secretion system-inactivated strain (ΔBscN). When hemolysis was induced, the culture turned red due to the release of hemoglobin. (B, C) Bar graphs show the results of hemoglobin measurement in the supernatant. Data show the average of the results of three triplicate experiments. *Statistically significant at *p* < 0.05.

Furthermore, to examine whether BcrH proteins were required for the function of the type III secretion system, L2 cells derived from rat lung epithelium were infected with the BcrH proteins-deficient strains. After infection, the amount of lactate dehydrogenase (LDH) released into the medium was analyzed to measure the efficiency of cell death induced by BteA, an effector, injected into mammalian cells via the type III secretion system (6, 10, 13). As a result, the amount of LDH released from mammalian cells was significantly reduced when infected with BcrH proteins-deficient strains compared to the wild-type strain (Fig. 4).

**Fig. 4.**
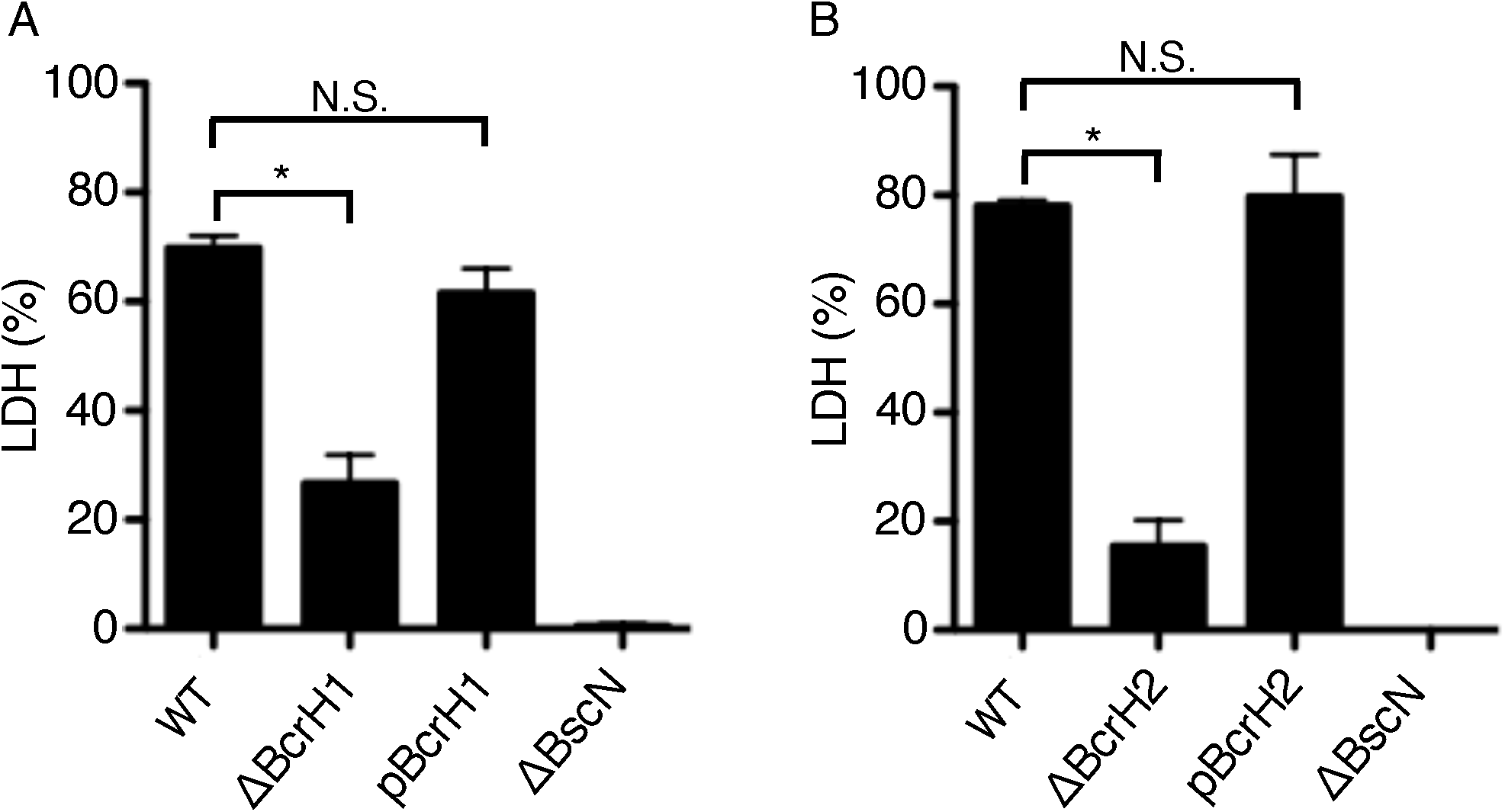
Analysis of the cytotoxicity of BcrH1- and BcrH2-deficient strains. The following *B. bronchiseptica* strains were analyzed: the wild-type strain (WT); the BcrH1-deficient strain (ΔBcrH1); the strain in which pBcrH1 was introduced into ΔBcrH1 (pBcrH1); the BcrH2-deficient strain (ΔBcrH2); the strain in which pBcrH2 was introduced into ΔBcrH2 (pBcrH2); and the type III secretion system-inactivated strain (ΔBscN). Each strain was cultured in SS medium and then infected into L2 cells at an MOI of 50 for 1 h. The results are shown as bar graphs in panels A and B. The amount of LDH in the medium was then measured. Data are the average from three triplicate experiments. *Statistically significant at *p* < 0.05; N.S., no significant difference.

These findings suggest that BcrH1 and BcrH2 are required for type III secretion system activity-dependent induction of hemolytic activity and cell death.

### BopB and BcrH1 form a complex within the *Bordetella* cytoplasm

In a previous study using immunoprecipitation assays with anti-BopB antibodies, BcrH2 was shown to form a complex with BopD and BopB in the bacterial cytoplasm (10). Accordingly, we here analyzed whether BcrH1 is included in the BopB-BopD complex. When anti-BcrH1 antibody was added to the lysate of the wild-type strain, the BopB signal was detected in the pellet fraction (Fig. 5A). However, in the immunoprecipitation assay using anti-BcrH1 antibody, we did not detect a clear signal of BopD in the pellet fraction (Fig. 5B). When immunoprecipitation was performed by adding BcrH2 antibody to the lysate of the wild-type strain, a faint BopB signal (Fig. 6A) and a clear BopD signal (Fig. 6B) were detected in the precipitate fraction. In contrast, no BopB signal was detected in the precipitate fraction from the ΔBopD lysate (Fig. 6A). These results suggest that BcrH1 forms a complex with BopB, and BcrH2 forms a complex with BopD, within the bacterial cell.

**Fig. 5.**
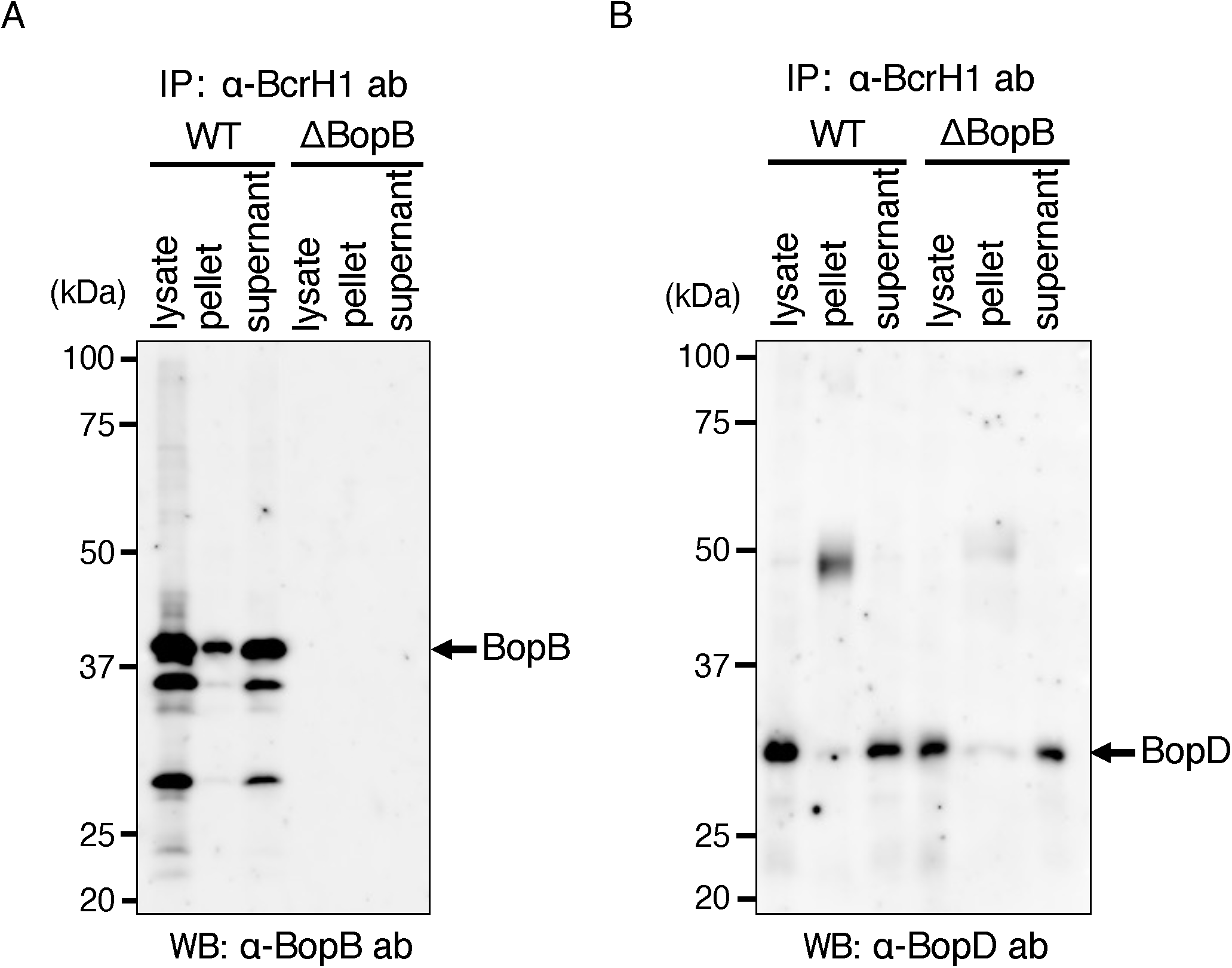
Analysis of the interaction between BcrH1 and pore-forming factors by immunoprecipitation. *B. bronchiseptica* cell lysates were prepared from the wild-type strain (WT) and the BopB-deficient strain (ΔBopB), and immunoprecipitation was performed using anti-BcrH1 antibody. Each sample was then separated by SDS-PAGE, and the results of Western blotting using anti-BopB antibody (A) and anti-BopD antibody (B) are shown.

**Fig. 6.**
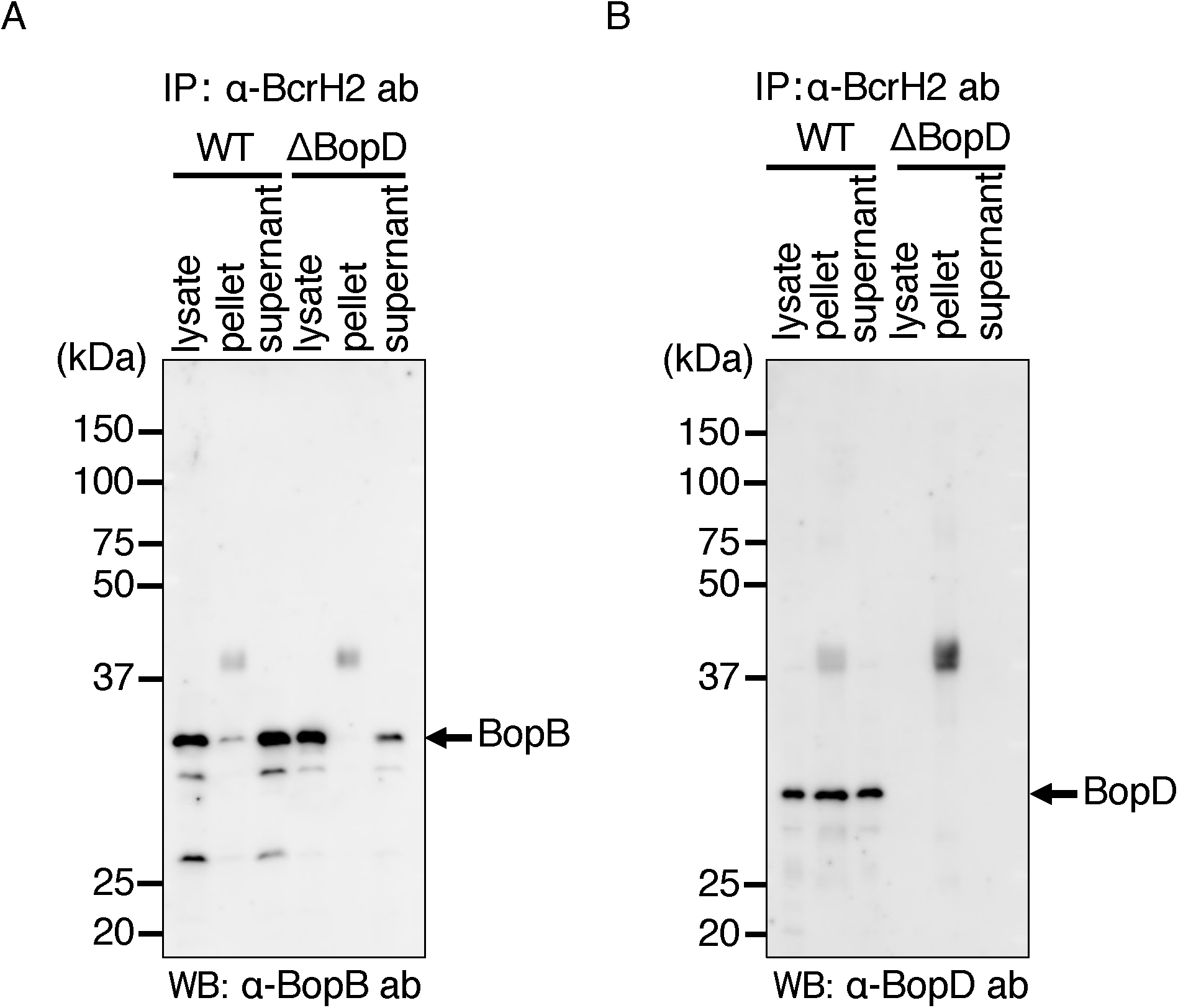
Analysis of the interaction between BcrH2 and pore-forming factors by immunoprecipitation. *B. bronchiseptica* cell lysates were prepared from the wild-type strain (WT) and the BopD-deficient strain (ΔBopD), and immunoprecipitation was performed using anti-BcrH2 antibody. Each sample was separated by SDS-PAGE, and the results of Western blotting using anti-BopB antibody (A) and anti-BopD antibody (B) are shown.

### BcrH1, but not BcrH2, is required for intracellular stability of BopB, and vice versa

Based on results in this study suggesting that BcrH1 and BcrH2 are chaperones for BopB and BopD, respectively, we next analyzed the specificity of these BcrH proteins for BopB and BopD. A double-deficient strain lacking both BcrH1 and BcrH2 (ΔH1H2) was constructed. Plasmids encoding the BcrH proteins were introduced into this double-deficient strain to construct strains ΔH1H2/pΒcrH1 and ΔH1H2/pΒcrH2, respectively. After culturing these strains in SS medium, whole cell fractions and supernatant fractions were prepared. These samples were developed by SDS-PAGE and then subjected to Western blotting with anti-BopB or anti-BopD antibodies.

We found no detectable BopB signal in the whole cell fraction of ΔH1H2 (Fig. 7), whereas a BopB signal was detected in the total cell fraction of ΔH1H2/pΒcrH1. In the case of BopD, no signal was detected in the whole cell fraction of ΔH1H2 (Fig. 7), but a signal was detected in the whole cell fraction of ΔH1H2/pΒcrH2. Furthermore, no BopD signal was detected in the total cell fraction of ΔH1H2/pΒcrH1, and no BopB signal was detected in the total cell fraction of ΔH1H2/pΒcrH2. These results strongly suggest that BcrH1 and BcrH2 are required for the intracellular stability of BopB and BopD, respectively.

**Fig. 7.**
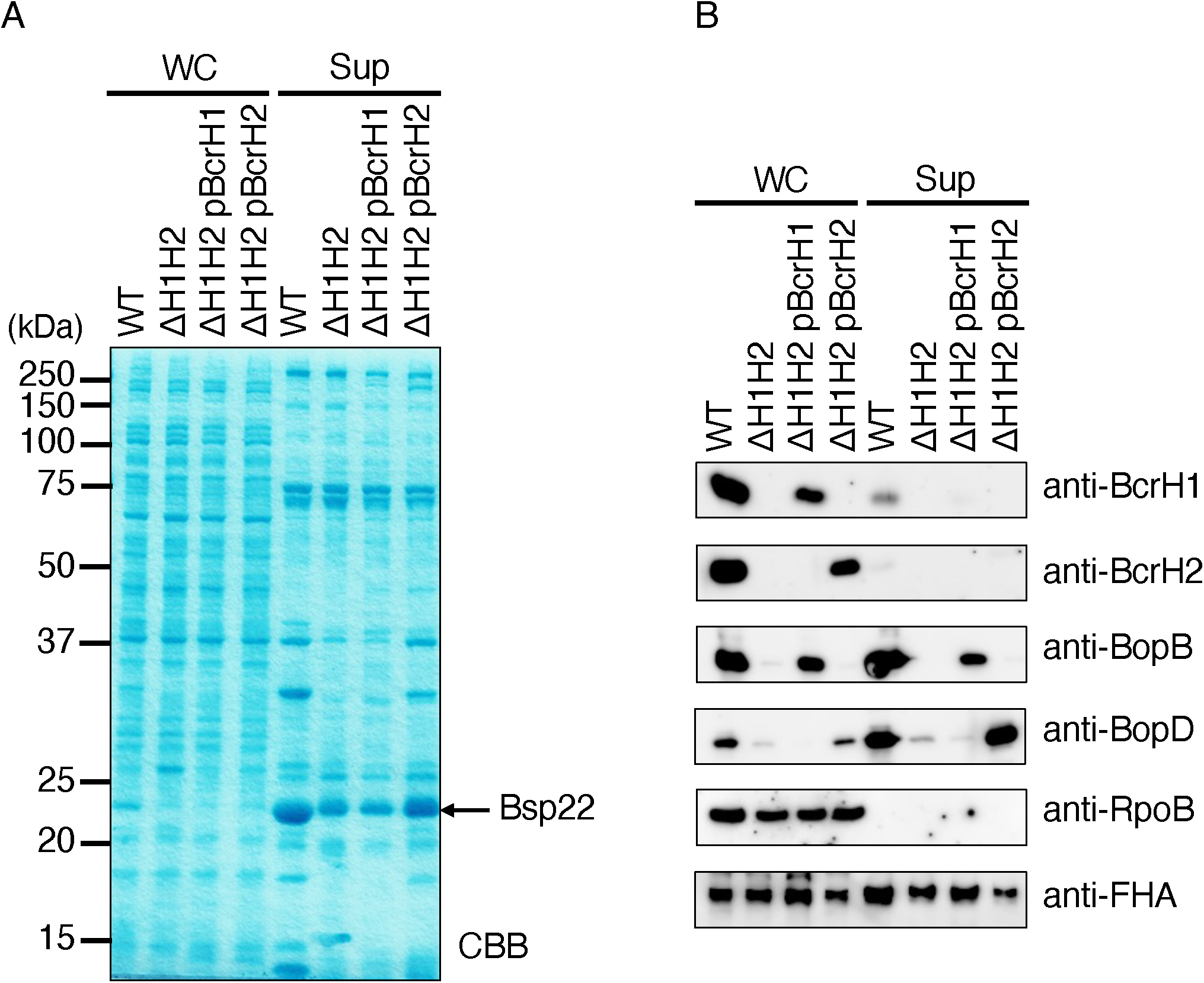
Analysis of production and secretion of type III secretory proteins in a BcrH1/BcrH2-double deficient strain. The following *B. bronchiseptica* strains were analyzed: the wild-type strain, a BcrH1/BcrH2 double-deficient strain (ΔH1H2), the strain with pBcrH1 introduced into ΔH1H2 (ΔH1H2 pBcrH1), and the strain with pBcrH2 introduced into ΔH1H2 (ΔH1H2 pBcrH2). Each strain was cultured in SS medium, and whole cell fractions (WC) and supernatant fractions (Sup) were prepared. (A) WC and Sup samples were separated by 12% SDS-PAGE and the gel was stained with CBB. Numbers indicate molecular weight markers in kDa. (B) WC and Sup samples were separated by SDS-PAGE, and then Western blotting was performed using anti-BcrH1 antibody, anti-BcrH2 antibody, anti-BopB antibody, anti-BopD antibody, RpoB (a control for cytoplasmic protein) antibody, and anti-FHA (a control for the type III secretion system-independent secreted protein) antibody.

## Discussion

In this study, we attempted to analyze the function of BcrH proteins to clarify their role in the infection process of *Bordetella*. Our results showed that the signal intensity of BopB and BopD in the whole cell fraction and the supernatant fraction of the BcrH1- and BcrH2-deficient strains, respectively, were significantly reduced compared to the wild-type strain (Figs. 1 and 2). The BcrH proteins were found to be required for the hemolytic activity (Fig. 3) and cytotoxicity (Fig. 4) of *B. bronchiseptica*. The immunoprecipitation experiments suggested that BcrH1 interacts with BopB and BcrH2 interacts with BopD (Figs. 5 and 6). The results of complementation of a BcrH1/BcrH2-double deficient strain (Fig. 7) strongly suggest that BcrH1 and BcrH2 are specific chaperones for BopB and BopD, respectively.

Western blotting using anti-BcrH1 antibody showed a weak BcrH1 signal in the supernatant fraction of the wild-type strain and the complemented strain (pBcrH1; Fig. 1B). Similarly, Western blotting using anti-BcrH2 antibody showed a weak BcrH2 signal in the supernatant fraction of the wild-type strain and the complemented strain (pBcrH2; Fig. 2B). In general, chaperones for type III secreted proteins are not secreted via the type III secretion apparatus after interacting with the type III secreted proteins in the bacterial cell. To investigate whether BcrH1 and BcrH2 are secreted via the type III secretion apparatus, we introduced pBcrH1 or pBcrH2 into the ΔBscN strain, which had lost type III secretion ability. We detected weak signals of BcrH1 and BcrH2 in the supernatant fractions prepared from these plasmid-introduced ΔBopN strains (data not shown), suggesting that BcrH1 and BcrH2 were not secreted from the type III secretion apparatus, but were nonspecifically eluted from the bacterial cells.

In immunoprecipitation experiments using anti-BcrH2 antibodies, a weak BopB signal was detected in the immunoprecipitated fraction of the wild-type strain (Fig. 6A). On the other hand, in immunoprecipitation experiments using the cell lysate of the BopD-deficient strain and anti-BcrH2 antibodies, no BopB signal was detected in the immunoprecipitated fraction (Fig. 6A). A weak BopB signal was detected in the immunoprecipitated fraction of the wild-type strain using anti-BcrH2 antibodies. Because BopB and BopD interact with each other (10), this result may indicate that BopB bound to BopD that had already interacted with BcrH2.

Many pathogenic bacteria that produce type III secretion systems form pores in the host cell membrane to translocate effectors into the host cell, and these pores are formed by pore-forming factors secreted by the type III secretion system (5). Numerous reports have shown that pore-forming factors have chaperones that play a role in maintaining their stability within the bacterial cell. *Pseudomonas aeruginosa* secretes PopB and PopD as pore-forming factors, and PcrH has been revealed as the chaperone for both PopB and PopD (14, 15). Similarly, SycD (LcrH) is a chaperone for the pore-forming factors YopB and YopD of *Yersinia pseudotuberculosis* (12), IpgC is a chaperone for the pore-forming factors IpaB and IpaC of *Shigella flexneri* (16), and SicA is a chaperone for the pore-forming factors SipB and SipC of *Salmonella enterica* serovar Typhimurium (17). In these bacteria, a single chaperone contributes to the stability of two pore-forming factors. *Bordetella* produces two proteins, BcrH1 and BcrH2, which have approximately 40% amino acid sequence similarity. A previous study reported by our group suggested that BcrH2 is a chaperone for BopD (10). However, it was unclear whether BcrH1 interacts with BopB by itself, or whether BcrH1 and BcrH2 contribute to the stability of BopD in a cooperative fashion. In this study, immunoprecipitation experiments showed that BcrH1 and BcrH2 interact specifically with BopB and BopD, respectively (Figs. 5 and 6). We also prepared a BcrH1/BcrH2 double-deficient strain and analyzed it by Western blotting using anti-BopB and anti-BopD antibodies (Fig. 7). When BcrH1 was complemented in the double-deficient strain, a BopB signal was detected, and when BcrH2 was complemented, a BopD signal was detected. These results strongly suggested that BcrH1 and BcrH2 function as specific chaperones for BopB and BopD, respectively (Fig. 7).

The reason why *Bordetella* produces two chaperones for pore-forming factor is unknown. There might be a *Bordetella*-specific regulation system associated with the type III secretion apparatus activity. The identification of other unknown proteins containing a BopB-BopD-BcrH1-BcrH2 complex in the *Bordetella* cytoplasm could provide clues for clarification of the regulation mechanism of the type III secretion system

## Materials and Methods

### Bacterial strains

The bacterial strains used in this study are shown in Table 1. *B. bronchiseptica* S798 was isolated from a pig (13). The BcrH2-deficient strain (ΔBcrH2) and BscN-deficient strain (ΔBscN) were created from *B. bronchiseptica* S798. BcrH2 and BscN were encoded at the *bsc* locus. *Escherichia coli* DH10B (Invitrogen) was used as a host for DNA cloning, and SM10λ*pir* (13) was used as a host for the suicide vector and for conjugative transfer.

### Bacterial culture

*B. bronchiseptica* was cultured on Bordet Gengou agar medium (18) for 2 days at 37°C, and single colonies were collected with a loop. These were suspended in Stainer-Scholte liquid medium (SS medium) (19). The bacterial suspension was adjusted to an absorbance of 0.23 at 600 nm, and cultured with shaking at 37°C for 18 h. Lysogeny Broth (LB) medium was used for *E. coli*. The antibiotics used for selection of strains were streptomycin at 30 μg/ml and ampicillin at 50 μg/ml.

### Gene-disrupted mutant creation

The plasmids used in this study are shown in Table 1, and the primers are shown in Table 2. To disrupt the *bcrH1* and *bcrH2* genes, PCR was performed using primers including each gene and its surrounding regions. Primers B1-bcrH1 and B2-bcrH1 were used for amplifying the *bcrH1* region, and primers B1-bcrH2 and B2-bcrH2 were used for amplifying the *bcrH2* region. The DNA fragments obtained by PCR were inserted into pDONR201 (Invitrogen) using the Gateway cloning system (Invitrogen) to obtain pDONR-bcrH1 and pDONR-bcrH2. To delete the internal sequence of each gene, inverse PCR was performed using the plasmid pDONR-bcrH1 or pDONR-bcrH2 as a template, and primer sets of R1-bcrH1 and R2-bcrH1 for bcrH1, or R1-bcrH2 and R2-bcrH2 for bcrH2. The PCR product was self-ligated using the In-Fusion Cloning System (Clontech) to create pDONR-ΔbcrH1 and pDONR-ΔbcrH2. These pDONR plasmids and the suicide vector pABB-CRS2 (20) were subjected to LR reaction to create pABB-CRS2-ΔbcrH1 and pABB-CRS2-ΔbcrH2. The pABB-CRS2 plasmids were introduced into *E. coli* SM10λ*pir*, and the resulting strains were mixed with a wild-type *B. bronchiseptica* to transfer the plasmids by conjugation as described previously (13). By ampicillin and sucrose selections, *B. bronchiseptica* ΔbcrH1 and ΔbcrH2 were obtained. To create a BcrH1/BcrH2 double-deficient strain, pABB-CRS2-ΔbcrH2 were transferred into *B. bronchiseptica* ΔbcrH1 and selected by antibiotics- and sucrose-containing plates as described above.

### Construction of expression plasmids for BcrH1 and BcrH2

PCR was performed using primers B1-bcrH1-comp and B2-bcrH1-comp with chromosomal DNA of the *B. bronchiseptica* S798 strain as a template to amplify the *bcrH1* gene. Similarly, primers B1-bcrH2-comp and B2-bcrH2-comp were used to amplify the *bcrH2* gene (Fig. 4). Each PCR product was cloned into vector pDONR-201 to obtain the plasmids pDONR201-bcrH1-comp and pDONR201-bcrH2-comp. Next, LR reactions were performed with a Multisite GATEWAY cloning system using the following: vector pRK-R4-R3-F (20), which can replicate in *Bordetella*; pDONR-fhaP (21), which encodes a promoter; pDONR-rrnB (21), which encodes a terminator; and pDONR201-bcrH1-comp or pDONR201-bcrH2-bcrH2. The reaction mixture was then used as the donor DNA in the transformation operation of *E. coli* DH10B. Plasmids were recovered from the transformants to obtain the pRK plasmids pBcrH1 and pBcrH2, which were then introduced into *B. bronchiseptica* strains by electroporation.

### Preparation of proteins from culture supernatants and whole bacterial cell lysates

Proteins secreted into bacterial culture supernatants and whole bacterial cell lysates were prepared as described previously (13). The loaded sample amounts were adjusted by the A_600_ of each bacterial culture in order to load samples prepared from the same amounts of bacteria. The protein samples were separated by SDS-PAGE and analyzed by Western blotting.

### SDS-PAGE and Western blot

The proteins were subjected to SDS-polyacrylamide gel electrophoresis (SDS-PAGE). Protein bands were detected by Coomassie brilliant blue (CBB) staining using one-step CBB staining solution (Biocraft). For Western blotting, the proteins in the SDS-PAGE gel were transferred to a PVDF membrane (Millipore). The membrane was immersed in Tris-buffered saline (TBS) containing 3% skim milk for 15 min for blocking, and then immersed in a primary antibody solution overnight at room temperature. The primary antibodies used were anti-BcrH2 antibody (13), anti-BopB antibody (13), anti-BopD antibody (10), anti-BopN antibody (22), anti-RpoB antibody (23), and anti-FHA antibody (24). To prepare the anti-BcrH1 antibodies, the peptides corresponding to the C-terminus region of BcrH1 (ERAESLRRSYARAD) were conjugated with haemocyanin from keyhole limpets (Sigma) by using 3-maleimidobenzoic acid N-hydroxysuccinimide ester (Sigma) respectively. These cross-linked peptides were used to immunize rabbits, and the resulting antisera were incubated with each peptide immobilized on epoxy-activated sepharose 6B (GE Healthcare) to obtain specific Ig-fractions. The PVDF membrane was then washed, and immersed in a solution containing horseradish peroxidase (HRP)-labeled Protein A solution (Sigma) or anti-mouse IgG solution (Sigma). Signal Enhancer HIKARI for Western Blotting and ELISA Solutions A and B (Nacalai Tesque) were used to dilute the antibody. Luminata Forte Western HRP Substrate (TaKaRa) was used to detect the signal.

### Immunoprecipitation

A 1-ml aliquot of overnight culture of *B. bronchiseptica* was centrifuged to obtain the bacterial pellet, which was then suspended in 1 ml of ice-cold PBS. The bacterial suspensions were sonicated using a Biorupter ultrasonicator (Cosmo Bio). The sonicated samples were centrifuged with 15,000 x g at 4°C for 15 min. The obtained supernatant was used as the bacterial lysate. Twenty-five microliters of anti-BcrH1 or anti-BcrH2 antibody was added to 180 μl of the bacterial lysate, and the mixture was stirred at low speed overnight at 4°C. Then, 30 μl of protein A sepharose (Sigma) was added, and the mixture was stirred at low speed for 1 h at 4°C. The beads were washed three times with 1 ml of ice-cold phosphate-buffered saline (PBS) containing 0.1% Triton, and then treated with 30 μl of 2×SDS-PAGE sample buffer and heated at 95°C for 3 min.

### LDH Assays

The *B. bronchiseptica* overnight culture was diluted with F-12K (Gibco). L2 cells were prepared in a 24-well plate. The infection assay was performed at a multiplicity of infection (MOI) of 50. After centrifugation at 1700 rpm for 5 min, the plates were incubated at 37°C and 5% CO2 for 1 h. The lactate dehydrogenase released from L2 cells was measured using a CytoTox96 Non-Radioactive Cytotoxicity Assay (Promega) according to the manufacturer’s protocol. Triton-X was added to the uninfected wells at a final concentration of 0.5%, and the LDH amount obtained from these wells was set as 100%. The relative value obtained from infected wells was calculated by subtracting the value of the uninfected well from the value of each infected well.

### Hemolysis Assay

Five volumes of PBS were added to rabbit red blood cells (Japan Biomaterials Center), and the mixture was centrifuged at 2800 rpm at room temperature for 5 min. The supernatant was removed, the precipitated red blood cells were suspended in an equal volume of PBS, and 50 μl of this red-blood-cell suspension was added to each well of a 96-well plate. *B. bronchiseptica* was cultured in SS liquid medium at 37°C for 18 h, and the bacteria in 2.5 ml of bacterial culture with an optical density at 600 nm (OD600) of 5.0 were collected by centrifugation and suspended in 500 μl of PBS. Bacterial suspensions were prepared based on the OD value of each culture so that an identical number of bacteria were infected with each strain. Fifty microliters of the bacterial suspension were added to each well containing red-blood-cell suspension. The 96-well plates were centrifuged at 1700 rpm at room temperature for 5 min, and then incubated at 37°C for 1 h in a 5% CO_2_ incubator. One hundred microliters of ice-cold PBS was added to each well, and the samples were mixed by pipetting. The plates were centrifuged with 2800 rpm at 4°C for 10 min, and 100 μl of the supernatant was collected and subjected to OD490 mesurement. As a control, fifty microliters of 10% Triton-X was added to the non-infected well, and the absorbance value obtained from these wells were set as 100%. The value obtained by subtracting the value of the non-infected well was used as a relative hemolytic activity.

### Statistical analyses

The statistical analyses were performed using the nonparametric test with a one-tailed *p-*value with Prism ver. 5.0f software (GraphPad, La Jolla, CA). Values of *p*<0.05 were considered significant.

## Acknowledgements

This work was supported in part by grants (23K06531 to A.K.) from the Ministry of Education, Culture, Sports, Science and Technology and the Japan Society for the Promotion of Science (KAKENHI), and a grant (JP24gm1610003 to A.K.) from the Japan Agency for Medical Research and Development (AMED). The funders had no role in the study design, data collection or analysis, the decision to publish, or the preparation of manuscript.

## References

1. Mattoo S, Cherry JD. 2005. Molecular pathogenesis, epidemiology, and clinical manifestations of respiratory infections due to Bordetella pertussis and other Bordetella subspecies. Clin Microbiol Rev 18:326–382.

2. Ahuja U, Shokeen B, Cheng N, Cho Y, Blum C, Coppola G, Miller JF. 2016. Differential regulation of type III secretion and virulence genes in Bordetella pertussis and Bordetella bronchiseptica by a secreted anti-σ factor. Proc Natl Acad Sci U S A 113:2341–2348.

3. Coburn B, Sekirov I, Finlay BB. 2007. Type III secretion systems and disease. Clin Microbiol Rev 20:535–549.

4. Büttner D. 2012. Protein export according to schedule: architecture, assembly, and regulation of type III secretion systems from plant- and animal-pathogenic bacteria. Microbiol Mol Biol Rev 76:262–310.

5. Cornelis GR, Van Gijsegem F. 2000. Assembly and function of type III secretory systems. Annu Rev Microbiol 54:735–774.

6. Kuwae A, Matsuzawa T, Ishikawa N, Abe H, Nonaka T, Fukuda H, Imajoh-Ohmi S, Abe A. 2006. BopC is a novel type III effector secreted by Bordetella bronchiseptica and has a critical role in type III-dependent necrotic cell death. J Biol Chem 281:6589–6600.

7. Panina EM, Mattoo S, Griffith N, Kozak NA, Yuk MH, Miller JF. 2005. A genome-wide screen identifies a Bordetella type III secretion effector and candidate effectors in other species. Mol Microbiol 58:267–279.

8. Blocker A, Gounon P, Larquet E, Niebuhr K, Cabiaux V, Parsot C, Sansonetti P. 1999. The tripartite type III secreton of Shigella flexneri inserts IpaB and IpaC into host membranes. J Cell Biol 147:683–693.

9. Neyt C, Cornelis GR. 1999. Insertion of a Yop translocation pore into the macrophage plasma membrane by Yersinia enterocolitica: requirement for translocators YopB and YopD, but not LcrG. Mol Microbiol 33:971–981.

10. Nogawa H, Kuwae A, Matsuzawa T, Abe A. 2004. The type III secreted protein BopD in Bordetella bronchiseptica is complexed with BopB for pore formation on the host plasma membrane. J Bacteriol 186:3806–3813.

11. Page AL, Parsot C. 2002. Chaperones of the type III secretion pathway: jacks of all trades. Mol Microbiol 46:1–11

12. Neyt C, Cornelis GR. 1999. Role of SycD, the chaperone of the Yersinia Yop translocators YopB and YopD. Mol Microbiol 31:143–156.

13. Kuwae A, Ohishi M, Watanabe M, Nagai M, Abe A. 2003. BopB is a type III secreted protein in Bordetella bronchiseptica and is required for cytotoxicity against cultured mammalian cells. Cell Microbiol 5:973–983

14. Discola KF, Förster A, Boulay F, Simorre JP, Attree I, Dessen A, Job V. 2014. Membrane and chaperone recognition by the major translocator protein PopB of the type III secretion system of Pseudomonas aeruginosa. J Biol Chem 289:3591–3601.

15. Job V, Matteï PJ, Lemaire D, Attree I, Dessen A. 2010. Structural basis of chaperone recognition of type III secretion system minor translocator proteins. J Biol Chem 285:23224–23232.

16. Page AL, Ohayon H, Sansonetti PJ, Parsot C. 1999. The secreted IpaB and IpaC invasins and their cytoplasmic chaperone IpgC are required for intercellular dissemination of Shigella flexneri. Cell Microbiol 1:183–193.

17. Tucker SC, Galán JE. 2000. Complex function for SicA, a Salmonella enterica serovar typhimurium type III secretion-associated chaperone. J Bacteriol 182:2262–2268.

18. Melton AR, Weiss AA. 1989. Environmental regulation of expression of virulence determinants in Bordetella pertussis. J Bacteriol 171:6206–6212.

19. Craig-Mylius KA, Stenson TH, Weiss AA. 2000. Mutations in the S1 subunit of pertussis toxin that affect secretion. Infect Immun 68:1276–1281.

20. Imaizumi A, Suzuki Y, Ono S, Sato H, Sato Y. 1983. Effect of heptakis (2,6-O-dimethyl) beta-cyclodextrin on the production of pertussis toxin by Bordetella pertussis. Infect Immun 41:1138–1143.

21. Yuk MH, Harvill ET, Cotter PA, Miller JF. 2000. Modulation of host immune responses, induction of apoptosis and inhibition of NF-kappaB activation by the Bordetella type III secretion system. Mol Microbiol 35:991–1004.

22. Nishimura R, Abe A, Sakuma Y, Kuwae A. 2018. Bordetella bronchiseptica Bcr4 antagonizes the negative regulatory function of BspR via its role in type III secretion. Microbiol Immunol 62:743–754.

23. Goto M, Abe A, Hanawa T, Suzuki M, Kuwae A. 2022. Bcr4 Is a Chaperone for the Inner Rod Protein in the Bordetella Type III Secretion System. Microbiol Spectr 10:e0144322.

24. Watanabe M, Nagai M, Funaishi K, Endoh M. 2000. Efficacy of chemically cross-linked antigens for acellular pertussis vaccine. Vaccine 19:1199–1203.

